# Development and Validation of Fluorescently Labeled, Functional Type I Collagen Molecules

**DOI:** 10.1101/2021.03.26.437209

**Authors:** Seyed Mohammad Siadat, Monica E. Susilo, Jeffrey A. Paten, Alexandra A. Silverman, Charles A. DiMarzio, Jeffrey W. Ruberti

## Abstract

While de novo collagen fibril formation is well-studied, there are few investigations into the growth and remodeling of extant fibrils, where molecular collagen incorporation into and erosion from the fibril surface must delicately balance during fibril growth and remodeling. Observing molecule/fibril interactions is difficult, requiring the tracking of molecular dynamics while, at the same time, minimizing the effect of the observation on fibril structure and assembly. To address the observation-interference problem, exogenous collagen molecules are tagged with small fluorophores and the fibrillogenesis kinetics of labeled collagen molecules as well as the structure and network morphology of assembled fibrils are quantified for the first time. While excessive labeling significantly disturbs fibrillogenesis kinetics and network morphology of assembled fibrils, adding less than ~1.2 labels preserves them. Applications of the functional, labeled collagen probe are demonstrated in both cellular and acellular systems. The functional, labelled collagen associates strongly with native fibrils and when added to an in vitro model of corneal stromal development, is endocytosed rapidly by cells and is translocated into synthesized matrix networks within 24 hours.

## 1. Introduction

Collagen is the most abundant protein in vertebrate animals. It is found in tissues that regularly experience tension, compression, and shear forces. The fibril forming types (I, II, III, V, XI, XXIV, and XXVII) are the principal load bearing molecules, with type I collagen being the most ubiquitous (e.g. the primary protein in blood vessel, bone, tendon, ligament, skin, sclera and cornea)^[1]^. Although connective tissue formation has been the subject of intense investigation for more than 100 years^[2]^, the cellular and molecular mechanisms that drive initial tissue formation and growth have yet to be clearly identified^[3]^. What is generally accepted is that resident mesenchymal cells are closely involved in the initial production of fibrils at or near their surfaces^[4]^. As neonatal tissue experiences rapid growth, cell density decreases^[5]^and cells have diminished access to directly influence individual fibrils^[6]^. Thus, we expect that cells may utilize at-a-distance techniques (e.g. surface energy modulation) to control collagen molecular incorporation and dissociation through chemical^[7]^or possibly mechanical “signals”^[8]^.

A remarkable feature of collagen fibrils is their dynamic remodeling in response to mechanical loading^[9]^which may have relevance to fatigue prevention under life-long cyclic loading^[10]^. Collagen molecules are constantly being synthesized, used to form fibrils, and removed under the control of a circadian clock to maintain collagen homeostasis^[11]^. However, the detailed mechanism by which the actual collagen molecules are added to (or subtracted from) the resident collagen fibrils is not known. To explore the possible mechanisms that govern fibril-molecule homeostasis at the fibril surface, it is important to be able to directly observe the motion of individual collagen monomers as they interact with extant, insoluble fibrils. Though collagen fibrils themselves are readily imaged via light (differential interference contrast (DIC)^[12]^; second harmonic generation (SHG)^[13]^), and electron microscopy^[14]^, dynamic tracking of their surface interaction with individual collagen monomers has not been performed. The effective diameter of collagen molecules are far below the Abbe diffraction limit of light and are thus undetectable via light microscopy, and electron microscopy requires a dehydrated, static sample^[15]^, preventing dynamic tracking.

Molecular labeling is a common strategy to enable light microscopy to detect structures below the wavelength of light. To detect formed collagen fibrils or adhered collagen molecules, collagen labeling is usually achieved by standard histological techniques which require cell fixation and therefore is not suitable for real-time imaging. Alternatively, more recent advancements in collagen labeling probes such as CNA35 – a collagen label based on a bacterial adhesion protein with specificity for collagen – permit time course visualization of collagen fibers without fixation^[16]^. CNA35 is small enough to diffuse into tissue and its binding to collagen is reversible. Therefore, it is a applicable probe for real-time, multi-color imaging^[17]^of newly formed fibrils and matrix development. Collagen molecules have been also labeled endogenously^[18]^and exogenously^[19]^with fluorescent tags to be detected during the fibrillogenesis process or during self-assembly. However, there is a concerning lack of concurrent investigations that examine the effect of the labeling procedure on fibrillogenesis kinetics of labeled collagen molecules as well as the structure and network morphology of assembled fibrils. Therefore, examination and quantification of collagen network dynamics with molecular scale resolution remains an important and unmet challenge.

Our goal was to fluorescently label and track collagen monomers, while minimizing the effect of labelling on the kinetics of collagen fibrillogenesis and on assembled fibril structure and network morphology. In this investigation, we use a small Alexa Fluor^™^ 488 (AF488) fluorescent dye (643 Da) to label single collagen molecules and quantify the functionality of labeled collagen molecules (AF488-Col). We then apply our <u>functional</u>, labeled collagen probe to 1) track molecular associations with native collagen fibrils in an acellular system and II) perform real-time imaging of extracellular matrix (ECM) development by corneal fibroblasts supplied with exogenous labelled collagen molecules. **Figure 1** provides an overview of the investigation.

**Figure 1.**
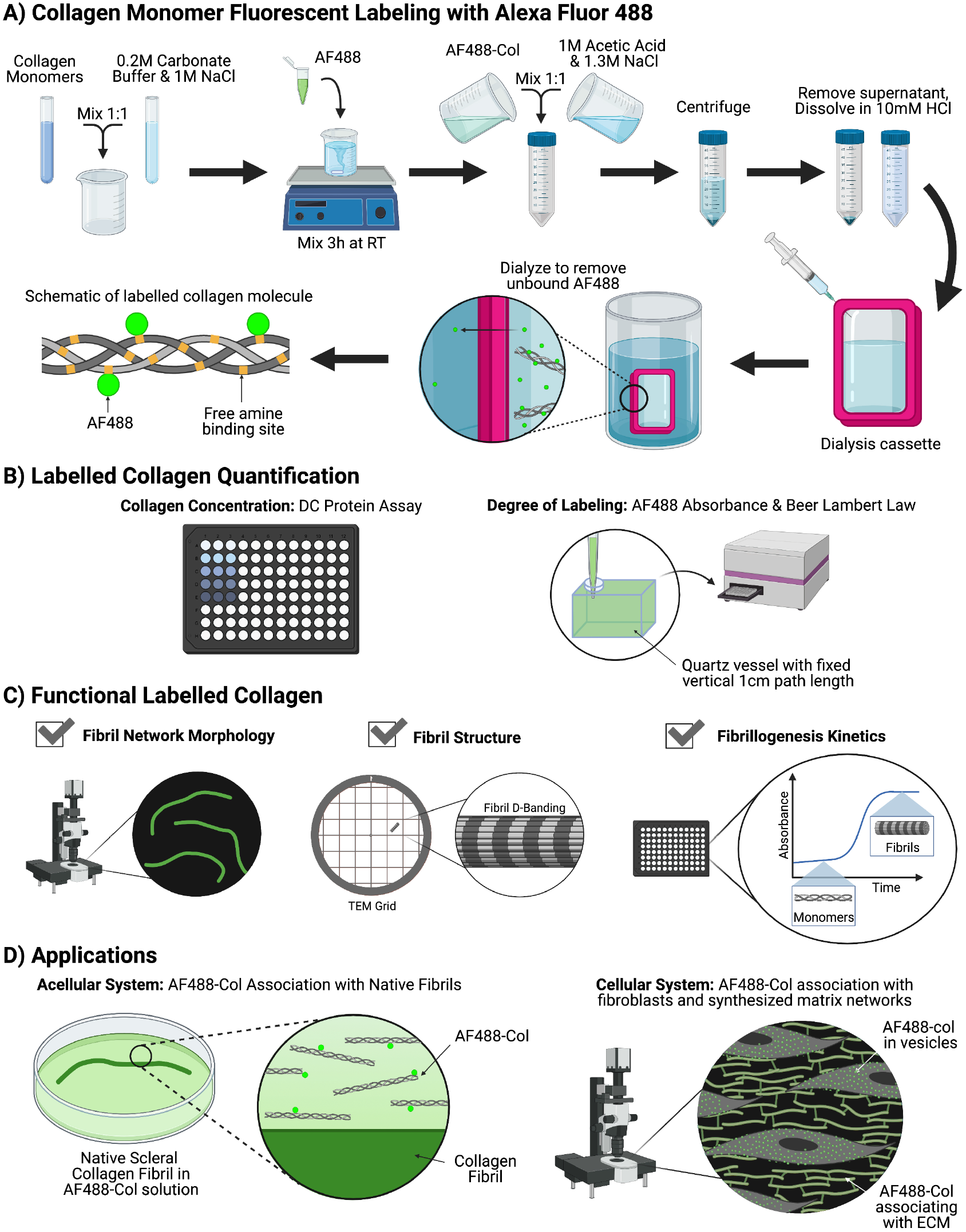
Study overview. A) Collagen molecules were labeled with AF488 fluorophores. B) Collagen concentration and DOL of AF488-Col were quantified using DC protein assay and Beer-Lambert law, respectively. C) Fibrillogenesis kinetics of AF488-Col as well as the structure and network morphology of assembled fibrils were quantified to confirm functionality of AF488-Col. D) Functional AF488-col was used in an acellular system were they associated with native scleral fibrils and in a cellular system were they associated with fibroblast cells and their synthesized ECM network.

## 2. Results and Discussion

### 2.1. Control of the Degree of Labelling (DOL)

Monomeric collagen was labelled with AF488 and the average DOL was reported as moles of dye per moles of collagen. **Figure S1** shows schematic of a labeled collagen molecule and **Figure S2** shows the effect of processing conditions (pH and molar ratio of AF488:collagen) on the final DOL.

### 2.2. Excessive Labeling Disturbs Fibrillogenesis Kinetics and Network Morphology

The first objective was to test the ability of our fluorescently labelled collagen molecules to form fibrils. To do this we took advantage of the well-characterized capacity of collagen molecules to spontaneously undergo de novo fibrillogenesis under physiological conditions^[20]^. The fibrillogenesis kinetics of AF488-Col was assessed via turbidity analysis. The sigmoidal turbidity curve observed during collagen assembly has two distinct phases: I) a lag phase during which there is no detectable change in absorbance and II) a linear growth phase during which the growth of fibrils absorbs light^[21]^. The fibrillogenesis kinetics of AF488-Col was studied and compared to unlabeled monomers (DOL of 0) by measuring the lag time, plateau time, and maximum change in absorbance at 313 nm.

**Figure 2** shows the fibrillogenesis kinetics for collagen solutions where monomers possessed a DOL of 0, ~0.5, ~0.8, ~1.2, ~2.1, ~4.1, and ~9.1. Fibrillogenesis was affected even at a relatively low DOL such as ~ 2.1 AF488:collagen, whereby the lag phase increased and the total absorbance decreased. Increasing DOL decreased the maximum absorbance (**Figure 2B**) and delayed the lag (**Figure 2C**) and plateau time (**Figure 2D**). Turbidity data analysis showed that fibrillogenesis kinetics were significantly changed for monomers labeled with an average DOL of 2.1 or higher. Fibrillogenesis of AF488-Col with DOL larger than ~9 did not show the typical sigmoidal shape turbidity curve. However, fibrillogenesis kinetics were preserved when monomers were labeled with a DOL of ~1.2 or less. **Table S1** summarizes the values of lag time, plateau time, and total absorbance for each tested condition.

**Figure 2.**
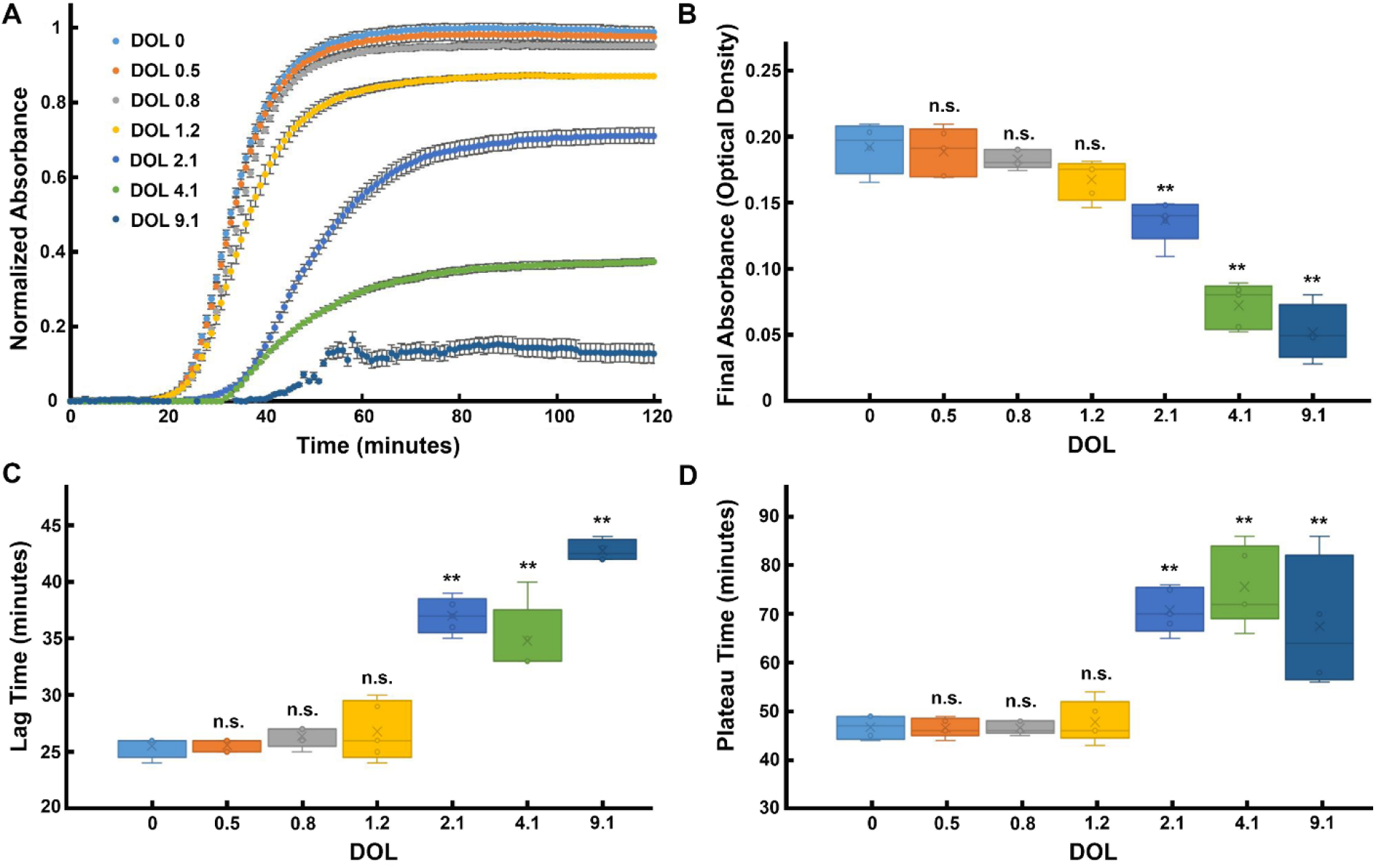
Fibrillogenesis kinetics of labeled collagen. (A) Turbidity curves of 50 μg/mL unlabeled (DOL 0) and AF488-Col at 37 °C. (B) Increasing DOL decreased the final absorbance. (C) Increasing DOL increased the lag time. (D) Increasing DOL increased the plateau time. Lag and plateau times were calculated as the time required to reach 10% and 90% of the maximum turbidity, respectively. Data are presented as mean ± standard deviation; n=4 or 5 replicates per group; Tukey’s honest significance test (** P<0.01; n.s., not significant).

The de novo fibril assembly data clearly demonstrate that the label interferes with the kinetics of assembly and that the effect decreases with decreasing the DOL. We then asked if the labels disrupted the morphology of the network or the nanostructure of the assembled fibrils. DIC, transmission electron microscopy (TEM), and fluorescence microscopy were used to determine if the fluorophores altered the fibrillar structure and network architecture. **Figure 3A** shows collagen fibrils formed from a 50 μg/mL solution of unlabeled collagen molecules. Monomers with DOL of up to ~2.1 (**Figure 3B**) produced an average fibril length and fibril count that was nearly indistinguishable from the unlabeled sample. Fibrils formed from collagen labeled with DOL of ~4.1 were both shorter and sparser in number (**Figure 3C**). When the labeling was increased to DOL of ~9.1, fibrillogenesis was strongly disrupted and non-fibrillar aggregates were mostly observed (**Figure 3D**). However, TEM images showed that the D-banding period – a hallmark of collagen fibril formation – remained unaffected (~67 nm) regardless of the DOL indicating that, despite the difference in network morphology, the labeling process did not impact the ultrastructure of the fibrils (**Figure 3E–H**). We thus concluded that while the label affected the kinetics of assembly at a high DOL (~2.1 or more), the final fibrillar structure remained “native”.

**Figure 3.**
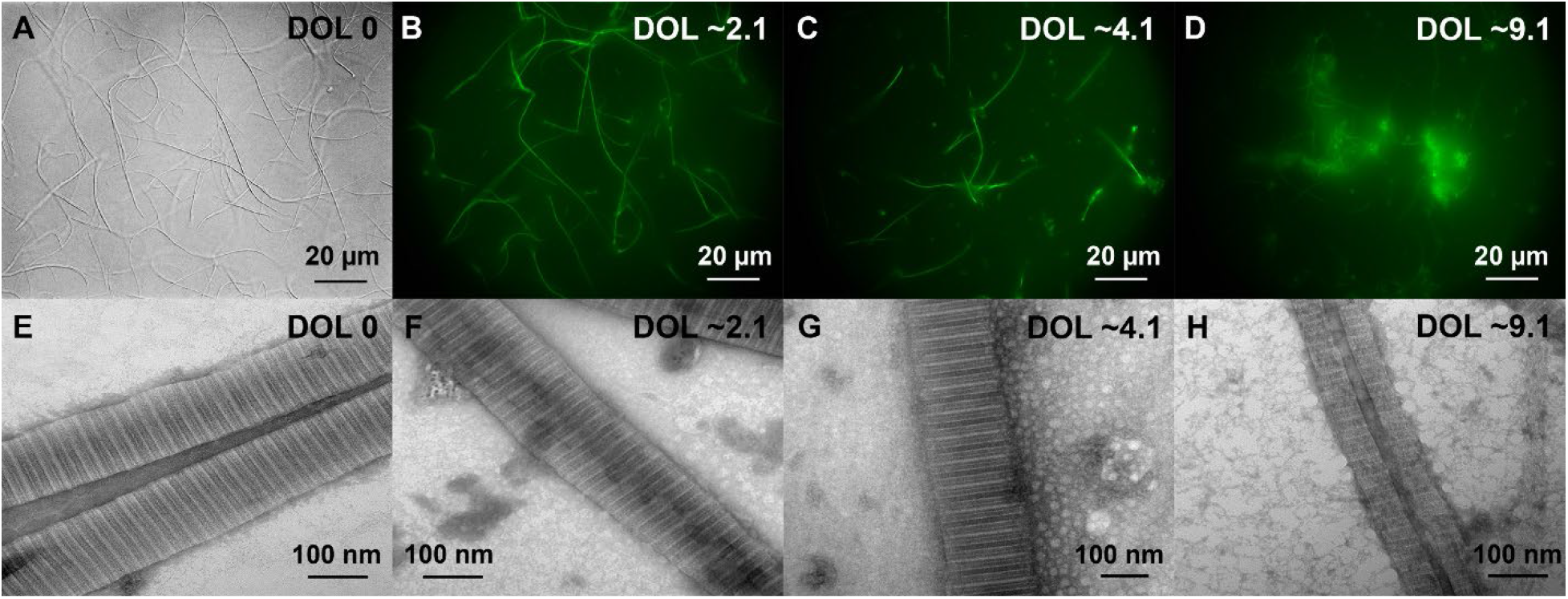
Collagen fibrils formed in neutralized solutions of 50 μg/ml collagen at 37°C. Top row shows DIC image of fibrils formed by unlabeled monomers (A) and fluorescent images of fibrils formed by AF488-Col with DOL of ~2.1 (B), ~4.1 (C), and ~9.1 (D). We used DIC imaging for the unlabeled fibrils because there was no fluorescence signal. Bottom row shows TEM images of fibrils formed by unlabeled monomers (E) and AF488-Col with DOL of ~2.1 (F), ~4.1 (G), and ~9.1 (H). While increasing the average DOL caused both shorter and sparser fibrils, the D-banding pattern remained unaffected (~67 nm).

We reported DOLs that were the average number of labels per collagen monomer and did not obtain information about the distribution of labels across the population of monomers. We expect that there will be some monomers that have no fluorophores, while some monomers have multiple fluorophores. Successful formation of fluorescent fibrils along with unchanged fibrillogenesis kinetics for monomers with low DOL (i.e. DOL of ~1.2 or less) suggests that monomers labeled with one fluorophore can assemble into fibrils with no significant change in the fibrillogenesis kinetics or D-banding. But monomers labeled with multiple fluorophores disrupt the fibrillogenesis kinetics and may not incorporate into fibrils. We suspect this is why the total absorbance (**Figure 2B**), is lower while the D-banding appears to be intact (**Figure 3E–H**).

### 2.3. Functional, Labeled Collagen Associates with Native Fibrils in an Acellular System

The next objective was to test the ability of our fluorescently labelled, functional collagen molecules to associate with native fibrils. Young, native collagen fibrils were trypsin extracted from 1-10-day old bovine sclera. Since proteoglycans can influence the association of collagen monomers with fibrils^[22]^, we determined if they were present on our extracted fibrils. The proteoglycan analysis showed that more than 99% of proteoglycans were removed from scleral fibrils during extraction (**Figure S3**). Thus, our system is testing the ability of collagen to interact with fibrils in the absence of proteoglycan control^[23]^(e.g. at the maximum association rate).

Proteoglycan-free scleral fibrils were placed in a perfusable, temperature controlled dish. Temperature was set at either 25 or 30 °C with a constant supply of 2 μg/mL functional AF488-Col (DOL of ~0.5). **Figure 4** shows that AF488-Col actively engage with the native fibril surface in a two-phase process (i.e. lag and growth) similar to that observed by turbidity during collagen fibrillogenesis de novo. Lag and plateau times in minutes (the times when the fibril intensity reached 10% and 90% of its maximum intensity, respectively) were measured as 59.9 ± 19.4 and 111.8 ± 37.4 at 25 °C (n=21 replicates) and 43.5 ± 11.7 and 97.8 ± 20.7 at 30 °C (n=23 replicates), respectively. The association rate and net accumulation of labeled monomers onto scleral fibrils were estimated as 2.8 ± 1.2 and 4.0 ± 1.5 molecules per *μ*m^2^ per minute and 114.3 ± 26.8 and 207.1 ± 55.4 molecules per μm^2^at 25 (21 replicates) and 30 °C (23 replicates), respectively. These estimations were based on measured fibril diameter (**Figure S4**) and fluorescent intensity of known AF488 concentrations (**Figure S5**). Given these estimations, at the plateau (saturation), the available space on the fibril surface would be 1/16 or 1/9 covered by exogenous monomer at 25°C and 30°C, respectively. Furthermore, using an Arrhenius plot, activation energy was measured as 12.6 kcal/mol.

**Figure 4.**
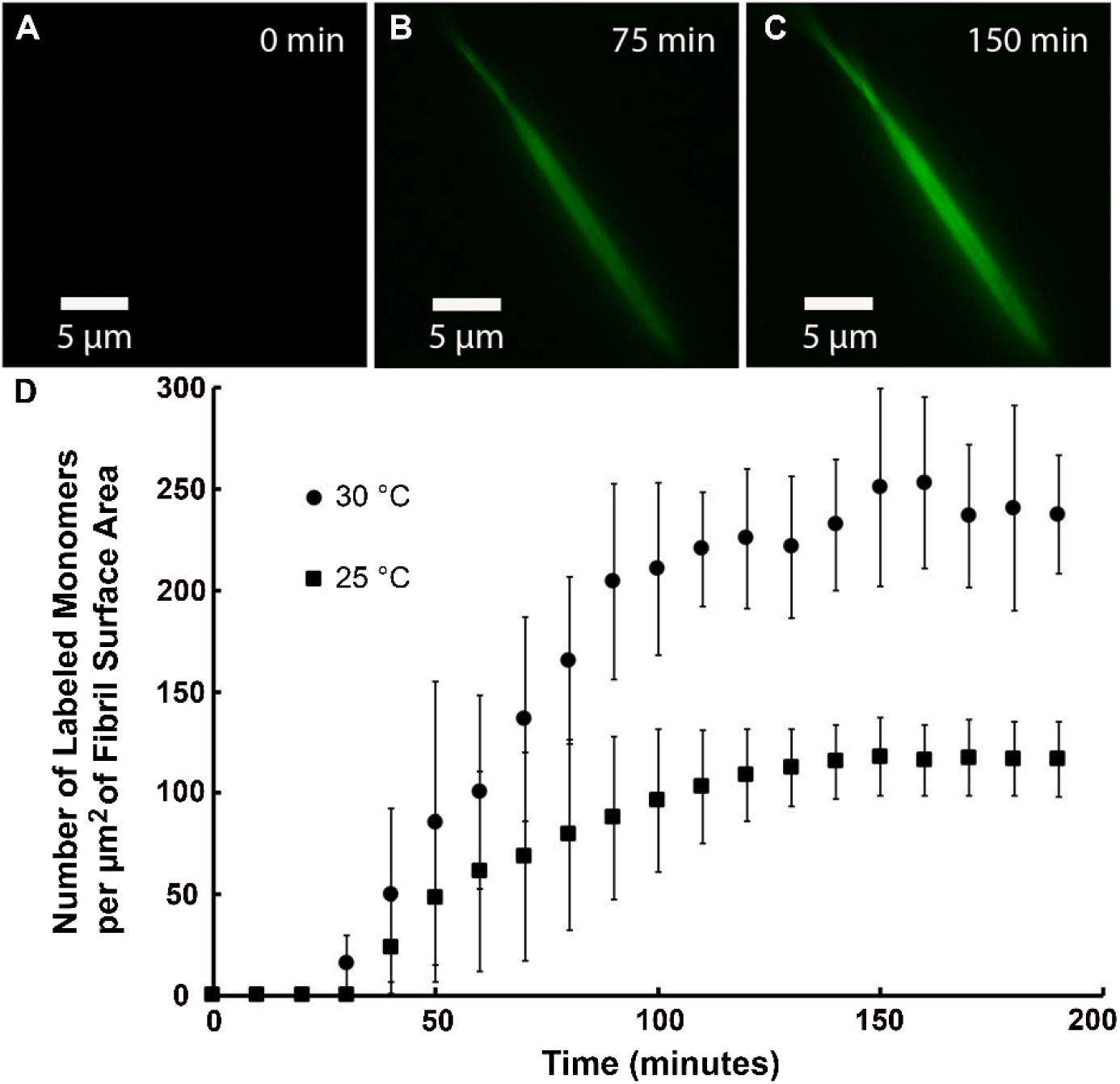
Association of functional AF488-Col with native fibrils. Exogenous labeled monomers associate with single, native sclera collagen fibrils with kinetics similar to fibril assembly. (A), (B), and (C) Fluorescent images show association of AF488-Col on a typical fibril over time at 25 °C. (D) The graph shows accumulation of AF488-Col onto sclera fibrils at 25 and 30 °C.

It has been shown that collagen molecules are produced at a relatively high rate by resident cells during tissue expansion^[24]^ and that the growing fibrils are bathed in a monomeric collagen solution that is constantly recycled^[25]^. We hypothesize that growing collagen fibrils are likely to be constantly exchanging molecules at their surfaces and that the surface exchange rates (k_on_ and k_off_) are modulated by both chemical and mechanical signals^[26]^. However, the baseline exchange rates for native collagen fibrils are not known. Part of the difficulty in establishing k_on_ and k_off_ is the insoluble nature of the collagen fibrils and the uncertainty of the nature of their association with soluble monomers. While estimation of the rate constants for collagen fibril surface erosion and adsorption requires relatively simple quantitative fluorescence microscopy, determining whether collagen monomers are incorporating functionally into extant fibrils is more difficult.

The activation energy of 12.6 kcal/mol calculated in the present study is lower than values reported in the literature for de novo collagen assembly in vitro (27 – 58 kcal/mol)^[27]^, suggesting that the activation energy for growth of a native fibril is less than formation and growth of a reconstituted fibril in vitro which one would expect. It has been shown that proteoglycans are removed from collagenous tissue by trypsin treatment^[23]^ similar to that which we employed. Therefore, the lateral growth of our scleral fibrils and the low activation energy might be attributed to the absence of proteoglycans. The scleral ECM has been shown to contain three major proteoglycans; biglycan, decorin, and aggrecan^[28]^. These proteoglycans can regulate matrix hydration^[29]^, interact with collagen fibrils^[30]^, and inhibit fibrillogenesis^[31]^. They have also been shown to control fibril diameter^[32]^by directly influencing molecular assembly^[22]^or lateral fusion of fibrils^[33]^.

During initial fibrillogenesis (i.e. **Figure 2**) the lag time we observe is likely due to the lack of pre-existing nucleation sites that must be first established. However, this is not the case in **Figure 4** where monomers are associating with extant native fibrils. In this experiment, where native sclera fibrils were incubated with AF488-Col, we used a constant supply of 2 μg/mL labeled monomers. At this subcritical concentration^[27a, 27c]^, monomers were unable to self-assemble at 25 and 30 °C at our experimental conditions. While monomers were below critical concentration and unable to form new fibrils in the solution, the native fibril could provide nucleation sites for formation of microfibrils. Therefore, the lag time we observe might be due to formation of microfibrils on the native fibrils.

The plateau region of the turbidity experiment (**Figure 2A**) is partially due to consumption of the reactant, while our monomer/fibril association experiment (**Figure 4**) had a constant supply of monomers. Our results suggest that at the plateau region of monomer/fibril association experiment, 6% and 11% of fibril surfaces were occupied by labeled monomers at 25 °C and 30 °C, respectively. Considering the constant supply of subcritical concentration of labeled monomers and saturation of scleral fibrils by different amounts of labeled monomers shows that in addition to proteoglycans, there must be other inhibiting factors that alter the availability of binding sites on fibrils and arrest the fibril’s growth in our system. However, we do not currently have a good explanation for the plateauing effect. Ultimately, we suspect that collagen fibrils reside in a state of dynamic equilibrium with the local extracellular milieu and that this equilibrium can potentially be altered by multiple factors (i.e. mechanical forces, matrix proteins, ionic concentration etc.) that could shift the delicate balance of the molecular association rates (k_on_ and k_off_) between fibrils and molecules.

### 2.4. Single Molecule, Multi-label Fluorescence Orientation Microscopy

We investigated the possibility of using <u>s</u>ingle molecule, <u>m</u>ulti-label fluorescence <u>o</u>rientation (SMO) microscopy to detect collagen monomers with 2 or more labels as they associate with collagen fibrils. The data we have provided in **Figure S6-8** and elsewhere[34] show that SMO microscopy has the potential to differentiate between monomers with one or multiple labels and that we can extract ellipses suggesting orientation of AF488-Col along the fibrils and ultimately determining whether collagen monomers are incorporating functionally into extant fibrils. However, more extensive work is necessary to fully determine the validity and limitations of SMO microscopy for these labelled collagen molecules.

### 2.5. Fibroblast Cells Incorporate Exogenously Labeled Monomers into Collagen Networks

The next objective was to test the functionality of exogenously labeled collagen monomers in a complex cellular system. Fibroblasts are the primary cell type in collagenous matrix^[35]^. They play an important role during fibril formation and remodeling conditions such as development^[36]^, fibrosis^[37]^, and wound healing^[38]^. In vitro, they can exert mechanical forces on collagen matrix^[39]^to translocate^[40]^, align^[41]^, and stabilize fibrils^[42]^. We added AF488-Col (DOL of ~0.9) to primary human corneal fibroblast (PHCF) cultures supplemented with ascorbic acid. **Figures 5** shows AF488-Col (in green) immediately after addition of labeled monomers to PHCF (**Figure 5A**), 24 hours after (**Figure 5B**), and 48 hours after (**Figure 5C**). Exogenously labeled collagen was internalized rapidly by PHCF and was translocated to the elaborated ECM networks within 24 hrs. Furthermore, when only AF488 was added to the cultured cells, it was taken up by PHCF but was not translocated to the ECM network suggesting that the collagen was the critical signal for transit to the matrix (**Figure 5D–F**).

**Figure 5.**
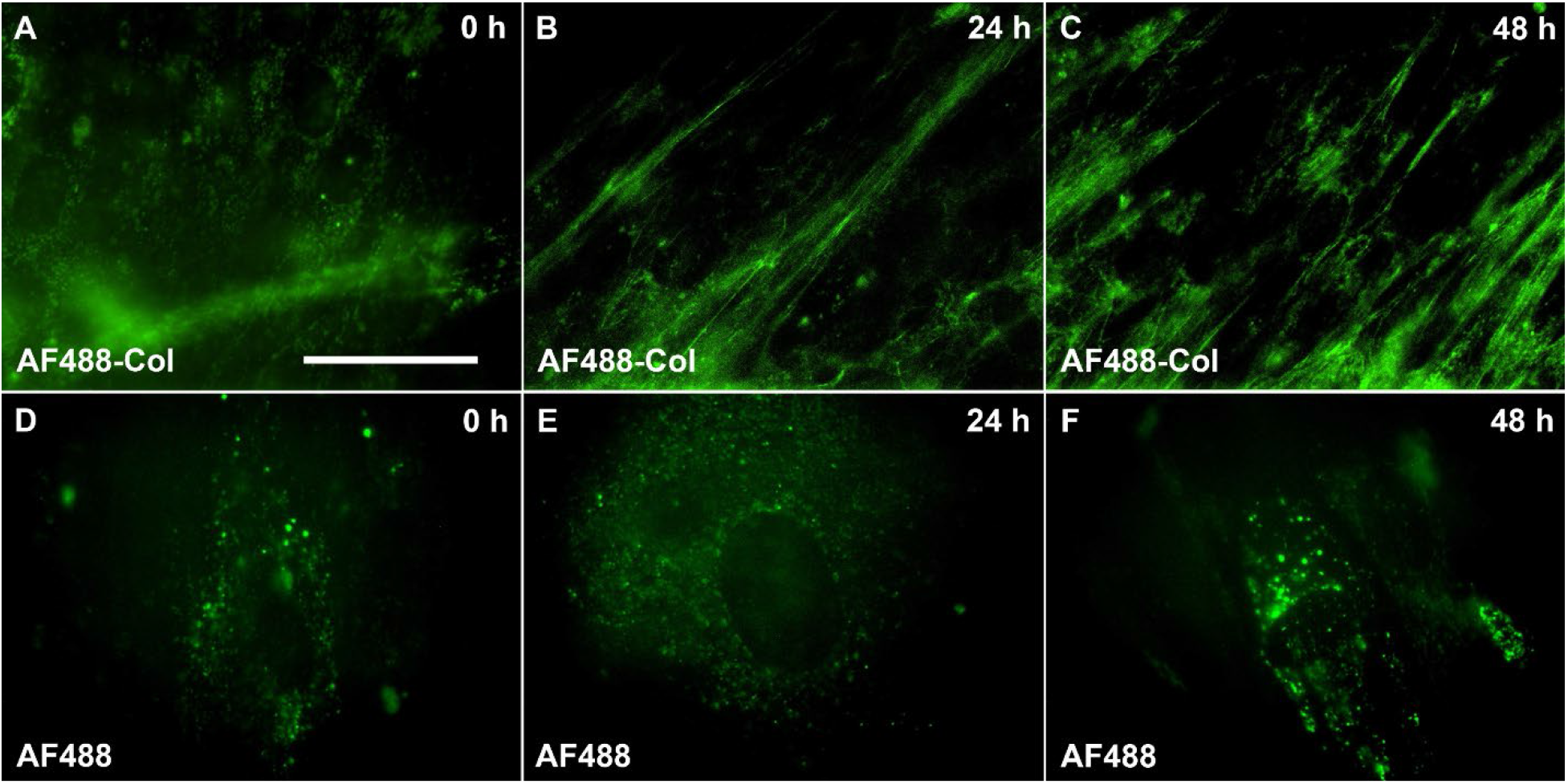
PHCF cells incorporate exogenously labeled monomers (green color) into an extracellular collagenous network. Top row shows the gradual formation of network after addition of labeled monomers (A), 24 hours after (B), and 48 hours after (C). Bottom row shows the control experiment where AF488 was added to collagen producing PHCF. AF488 did not associate with fibrillar ECM immediately after addition (D), 24 hours after (E), and 48 hours after (F). Scale bar is 50 μm and images are at the same magnification.

Our expectation was that exogenous collagen would “home” to fibrils and incorporate into them directly. However, we uncovered a very interesting phenomenon. Currently, the literature proposes a number of mechanisms that operate during early tissue formation, organization and growth. For instance, it has been suggested that fibrils are formed intracellularly^[43]^, extracellularly^[8b]^, or that fibril growth is the result of linear and lateral fusion of immature intermediate fibrils^[44]^. We are thus keenly interested in the relationship between the young fibril surface which is not yet fully cross-linked and the free collagen monomers which reside in the adjacent extracellular solution^[25]^. Using functional AF488-Col <u>with unaltered fibrillogenesis kinetics</u> we showed that our labeled monomers actively engage with native fibrils in an acellular system and interact with an ECM network elaborated under cellular control. Such a tool will enable us to ask fundamental and quantitative questions about the dynamics of molecular assembly and disassembly at the collagen fibril surface.

## 3. Conclusions

We have demonstrated a potentially powerful tool to not only visualize and track collagen molecules in real-time, but it will also permit investigation of fibrillogenesis kinetics and collagen network dynamics in vitro with molecular scale resolution. We showed that it is critical to perform characterization experiments to ensure that labelling does not alter fibrillogenesis kinetics or fibril morphology, particularly if one is investigating collagen assembly in real time. We were fortunate to discover that at low DOL (i.e. less than ~1.2 fluorophore per monomer), our labeling strategy had no detectable effect on collagen assembly kinetics, fibril morphology, or D-banding. However, increasing the DOL to more than ~2.1 fluorophore per monomer, disrupted the fibrillogenesis kinetics, but preserved D-banding periodicity. Our data indicate that it is not sufficient to only examine the D-banding of fibrils formed by labeled monomers. Fibrillogenesis kinetics could still be severely disrupted even when the fibrils look native.

We have confirmed here that our functional, labeled collagen has promising applications and is capable of dynamically tracking collagen monomers interacting with 1) single collagen fibrils under acellular conditions of interest and 2) an ECM network elaborated under cellular control. To show feasibility for the latter application, we applied our functional, labelled collagen to an in vitro model of corneal stromal development. We demonstrated that fibroblast cells use our exogenously labeled monomers to form collagen networks in vitro. The next step will be to use our fluorescently labeled collagen monomers in models of collagenous tissue growth and remodeling to further understand collagen monomer/fibril biophysics.

## 4. Experimental Section

### 4.1. Collagen Labeling

Bovine collagen solution (5026-50ML, TeloCol, Advanced BioMatrix) was diluted to 1 mg/ml using 10 mм hydrochloric acid (SA56-1, Fisher Scientific), and then mixed 1:1 with 0.2 M sodium carbonate–bicarbonate buffer (24095, Polysciences) containing 1 M sodium chloride (S671-3, Fisher Scientific). The labelling protocol was carried out at pH 7.5, 8.0, and 8.5 to probe the effect of pH on efficiency of labeling. Lysine residues are most efficiently labeled at pH values of 8.5-9.5; however, the lower pH range was selected to increase the probability of attaching one fluorophore to the *N*-terminal^[45]^and improve the average distance between fluorophores on each collagen molecule. Next, AF488 (A37570, Alexa Fluor 488 TFP ester, Invitrogen) was diluted to 0.5 mg/mL using dimethyl sulfoxide (D12345, Life Technology) and slowly added to collagen solution in 1x to 15x molar excess. The labeling reaction was carried at room temperature and gently stirred for 3 hours.

### 4.2. Purifying the Labeled Collagen

AF488-Col was mixed 1:1 with 1 M acetic acid (AC124040025, Acros Organics) containing 1.3 M sodium chloride to precipitate the collagen^[46]^. Next, the sample was centrifuged for 1 hour at 10000×g, the supernatant was removed, and the pellet was dissolved in 10 mM hydrochloric acid for 12 hours at 4 °C. AF488-Col was injected into a 3.5 kDa MWCO dialysis cassette (66110, Thermo Scientific) and dialyzed for 3 days against 100 times the volume of 10 mM hydrochloric acid at 4 °C to remove any unbound dye from the protein solution. To ensure the efficiency of dialysis process, AF488 (without collagen) was injected into the dialysis cassettes and dialyzed under the same conditions as AF488-Col. After 3 days in the dialysis cassettes, the samples’ absorbance was measured at 494 nm using a 1 cm pathlength cuvette and a plate reader spectrophotometer (PowerWave XS, BioTek Instruments). No free fluorophore was detected in the cassettes that had only AF488 (without collagen), indicating that all fluorophores remained in AF488-Col samples were bound to collagen molecules.

### 4.3. Determining Collagen Concentration and Degree of Labeling

The collagen concentration of AF488-Col was determined using DC Protein Assay (500-0114, Bio-Rad Laboratories). AF488 concentration was calculated based on the Beer-Lambert law:

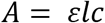

where *A* is the absorbance, *ɛ* is the molar extinction coefficient, *l* is the optical path length, and *c* is the concentration. The AF488-Col absorbance was measured at 494 nm in a 1 cm pathlength cuvette. A cuvette with long and calibrated pathlength was used to improve measurement accuracy and repeatability. Absorbance of the cuvette filled with 10 mM HCl was used for background absorbance. It is extremely important to properly clean the cuvette before each use and measure the background absorbance to ensure repeatability. The cuvette was filled gently using 100 μL pipette tips to avoid any bubble formation. The results were validated using standard samples with known concentration of AF488 (**Figure S9**). The degree of labeling was calculated as moles of dye per moles of collagen.

### 4.4. Spectrophotometry

To quantify the fibrillogenesis kinetics of each batch of AF488-Col solution, a plate reader spectrophotometer (PowerWave XS, BioTek Instruments) was used to measure the turbidity curve associated with assembly. This was compared against unlabeled solution at the same concentration to determine the impact of labeling. Collagen solutions were prepared at 50 μg/mL in phosphate buffered saline (PBS), pH 7.3, and room temperature. The spectrophotometer temperature was set at 37 °C. 100 μL samples were held in a 96 well plate and absorbance of the collagen solutions were read every 30 seconds at 313 nm. The lag and plateau times were calculated respectively as the time when the turbidity curve reached 10% and 90% of the maximum absorbance.

### 4.5. Scleral Fibril Extraction

Native collagen fibrils were trypsin extracted^[47]^from 1-10 day old bovine sclera. Briefly, the scleral tissue was isolated from the cornea, optic nerve, muscle, and surrounding adipose tissue. The vitreous humor was removed and the retina, tapetum lucidum, and choroid layers were debrided from the sclera, and the sclera was then cut into smaller pieces (approximately 1 cm by 2 cm). Some shallow cuts were made on each piece to increase surface area for fibril extraction. Each piece of sclera was rinsed with PBS and then placed into 20 mL of pH 7.5-8.0 extraction medium containing: >1000 BAEE unit/mL trypsin (9002-07-7, Sigma-Aldrich), 4 mM ethylenediaminetraacetic acid (60-00-4, Sigma-Aldrich), 0.1 M tris-HCl (1185-53-1, Sigma-Aldrich), and 0.05% sodium azide (26628-22-8, Sigma-Aldrich). Samples were shaken for ~5 minutes until the sclera swelled and some fibrils were removed. Fibril suspensions were centrifuged at 5000×g for 30 minutes. The supernatant was removed, and the fibril pellet was resuspended and stored in PBS containing 0.05% sodium azide at 4 °C.

### 4.6. Proteoglycan Quantification

Fibril suspensions in trypsin extraction medium were centrifuged for 30 minutes at 14,500×g and the pellet was suspended overnight in 300 μL of 10 mM HCl at 4°C. 50 μL of 8 M GuHCl solution (Acros Organics, 50-01-1) was added to each sample and incubated overnight at 4°C. Then, 50 μL of SAT solution (used to dilute alcian blue solution) containing 30 μL of 18 M sulfuric acid (Acros Organics, 7664-93-9), 75 μL of triton x-100 (Sigma-Aldrich, 9002-93-1), and 10 mL water was added to each sample, vortexed, and incubated at room temperature for 5 min. Alcian blue solution containing 20 μL of 18 M sulfuric acid (Acros Organics, 7664-93-9), 1 mL of 8 M GuHCl, 10 mL of 1% alcian blue in 3% acetic acid (Sigma-Aldrich, B8438), and 10 mL water was prepared. 750 μL of alcian blue working solution (5 SAT: 9 water: 1 alcian blue solution) was added to each sample and incubated for 1 hour at 4°C. Samples were centrifuged for 15 min at 12,000×g and supernatant was discarded. Pellets were suspended in 500 μL washing solution containing 8mL dimethyl sulfoxide (Fisher Scientific, 67-68-5), 0.2 g magnesium chloride hexahydrate (MP Biomedicals, 67-68-5), and 12 mL water. Samples were centrifuged for 15 min at 12,000×g and supernatant was discarded. Pellets were suspended in 500 μL of disassociation solution containing 6.6 mL of 8 M GuHCl, 3.3 mL of 2-propanol (Fisher Scientific, 67-63-0), and 25 μL of triton X-100 (Sigma-Aldrich, 9002-93-1). Absorbance of samples were read at 600 nm and concentration of proteoglycans were measured against a standard concentration of chondroitin sulfate (Sigma-Aldrich, 39455-18-0).

### 4.7. Imaging

DIC and fluorescent images were taken on a Nikon inverted microscope (ECLIPSE TE2000-E) equipped with a 60X oil objective (Nikon’s CFI Apochromat TIRF Series, Numerical Aperture: 1.45), a CoolSNAP EZ CCD Camera, and a high Speed EMCCD Camera (iXon Ultra 897, ANDOR) for detecting single molecules.

### 4.8. Fibril Diameter Measurement using TEM and DIC

Since DIC microscopy cannot directly measure submicron fibril diameters, a correlative method^[12]^was applied. Fibril diameters ranging between 60-240 nm can be determined using DIC and a 60x objective (NA 1.45), if the fibril diameter distribution and the DIC edge intensity shift (DIC-EIS) distribution are known for a given batch of fibrils^[48]^. Therefore, the diameter distribution was obtained from TEM images (**Figure S4A**). To prepare samples for TEM, 20 μL of sclera suspension was added on a 300 mesh formvar coated TEM grids (01701-F, Tedpella). After 10 minutes, when fibrils attached on the grid, the grid was wicked dry using a filter paper and stained 3 times on a droplet of 1.5% uranyl acetate for 3 seconds each time. The grid was dried with filter paper and imaged with a JEOL JEM 1010 transmission electron microscope.

To obtain DIC-EIS distribution of the same batch of sclera fibrils, the fibril suspension was pipetted onto a coverslip coated with 1% bovine serum albumin (BSA, A2153-10G, Sigma Aldrich) and given 1 hour to adhere. Next, only the fibrils oriented perpendicular to the shear axis of the light path were selected for intensity measurements to maximize signal strength and minimize error (**Figure S4B**). A z-scan was used to find the maximum DIC-EIS (**Figure S4C**). Then DIC-EIS measurements were taken along the middle section of fibrils using a custom MATLAB code. The average DIC-EIS was determined for 88 fibrils to establish the intensity distribution and correlated to the diameter of 800 fibrils measured with TEM.

### 4.9. AF488-Col Association with Native Scleral Fibrils Experiment

A glass bottom, ITO coated Delta-T dish (04200417C, Bioptechs) and a perfusable coverglass lid (0420031216, Bioptechs) were plasma cleaned and coated with 1% BSA at 4 °C overnight. The dish was placed on the microscope stage and 1 mL of sclera fibril suspension was added. After an hour incubation for fibril attachment, 20 mL of 1X PBS was perfused into the dish at 20 mL/h using a syringe pump. A temperature controller (5410429, Bioptechs) was set at either 25 or 30 °C, and the fibrils were imaged prior to the addition of labeled monomers. Next 400 μL of 10 μg/mL AF488-Col (DOL of ~0.5) was added to the dish at 12 mL/h for 2 minutes to rapidly adjust the collagen concentration inside the dish to 2 μg/mL (perfect mixing was assumed). Then 5 mL of 2 μg/mL AF488-Col was added at 1 mL/h to maintain constant concentration inside the chamber. The collagen concentration was kept at 2 μg/mL (sub-threshold for new fibril formation) to prevent self-assembly^[27c, 49]^and formation of new fibrils independent of the native fibril in the dish. DIC and fluorescent images were taken every 10 minutes for 3 hours. As a control experiment, AF488-Col was added to the dish in the absence of native scleral fibrils to ensure that collagen concentration was at sub-threshold for new fibril formation.

### 4.10. Image Analysis of Fluorescent Images

Incorporation rate and total accumulation of AF488-Col into the scleral fibrils were measured by tracking the fluorescent intensity of a region of interest (ROI) defined along the middle section of fibrils. Standard samples of AF488-Col with known collagen and AF488 concentrations were used to correlate fluorescent intensity of the ROI to number of collagen molecules (**Figure S5**). For each fibril, a 32 μm by 100 μm ROI was defined and the background-subtracted fluorescent intensity along the fibril length was correlated to the number of collagen molecules^[50]^within the effective depth of field.

Depth of field (*d*_*f*_) was calculated as sum of the wave and geometrical optical depth of field:

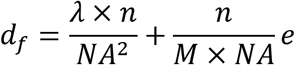

where *λ* is the wavelength of light, *n* is the refractive index of the immersion oil, *NA* is the objective numerical aperture, *M* is the objective lateral magnification, and *e* is the camera pixel size^[51]^.

### 4.11. Arrhenius Plot and Activation Energy Measurement

The reaction rate of AF488-Col binding to the surface of a sclera fibril was measured as the increase of fluorescent intensity with time. The reaction rate constant (*k*) was then calculated using the reaction rate equation:

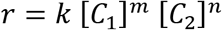

where *r* is the reaction rate and *[C_1_]* and *[C_2_]* are the collagen concentration in the solution and on the fibril surface, respectively. The surface concentration was calculated under the assumption that the monomers were packed with a lateral intramolecular Bragg spacing of 1.58 nm^[52]^ and a periodic gap region of 36 nm between the *N*-terminus of one molecule and the *C*-terminus of the next molecule^[53]^. Therefore, *[C_2_]* was calculated as 3.12×10^−9^ mol/m^2^ and *[C_1_]* was set at 2 μg/mL. The reaction orders, *m* and *n*, have been shown to be first order during in vitro collagen fibrillogenesis^[27b]^. The reaction rate constant was determined at 25 and 30 °C and the Arrhenius equation:

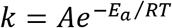

was used to determine both the activation energy (*E_a_*) and the pre-exponential factor (*A*) where *T* is temperature in Kelvin and *R* is the universal gas constant.

### 4.12. Potential Detection of Multiply-Labeled Collagen Molecules’ Orientation

When collagen is labeled with two or more fluorophores separated by a large enough distance, it is possible to fit an ellipse to the pixel representation. Thus, we can differentiate between singly-labeled and multiply-labeled monomers (**Figure S6-8**). We asked if we could determine whether it is possible to detect multiply labeled monomers (2 or more label on each monomer) as they associate with single collagen fibrils <u>and</u> detect the direction of the AF488-Col.

Labeled and unlabeled monomers were mixed 1:20000 in PBS at pH 7.3 and a total collagen concentration of 50 μg/mL. The mixture was incubated at 37 °C in a Delta-T dish to allow for fibrillogenesis on the glass surface. After reaching equilibrium, reconstituted fibrils and fluorescently labeled collagen monomers were imaged at 60x with high speed, highly-sensitive EMCCD camera (iXon Ultra 897, ANDOR). The image processing to obtain the orientation of the ellipse major axis was performed in Matlab (Mathworks, Inc., Natick MA) as follows: images were uploaded into Matlab, filtered using a Gaussian filter, and then background subtracted. The binary image was then processed using the function regionprops to obtain the orientation of the ellipse.

### 4.13. Cell Culture

PHCF cultures were established from 74 year donor corneas as described in Bueno et al.^[54]^. PHCF were seeded on temperature-controlled Delta-T dishes at passage 3-5 at a concentration of 30,000 cells/cm^2^. Once confluent on day 3, 0.5 mM *L*-ascorbic acid was added to stabilize collagen produced by the culture, and to permit the culture to construct a 3-D matrix of multiple orthogonal cell layers. On day 6, 1.5 mL of CO_2_ infused DMEM containing 10% fetal bovine serum, 1% antibiotic/antimycotic, 0.5 mM *L*-ascorbic acid and 1 μg/mL AF488-Col with a (DOL of ~0.9) was added to the cells. The culture was imaged using high-resolution 60x objective at 37°C by taking a photo immediately after AF488-Col addition, 24 hours after, and 48 hours after. The media (CO2 infused DMEM containing 10% fetal bovine serum, 1% antibiotic/antimycotic, 0.5 mM *L*-ascorbic acid) was exchanged before imaging at 24 and 48 hours. In a control experiment, only AF488 (at the same concentration) instead of AF488-Col was added to the cell culture.

### 4.14. Statistical Information

Data are shown as mean ± standard deviation. The sample number (*n*) indicates the number of independent biological samples in each experiment. Differences in means were considered statistically significant at *P* < 0.05. Significance levels are: * *P* < 0.05; ** *P* < 0.01; *** *P* < 0.001; **** *P* < 0.0001; *n.s.*, not significant. The sample sizes and the results of hypothesis tests are reported for each experiment in the results section.

## Supporting Information

Supporting Information is available from the Wiley Online Library or from the author.

## Authorship Contribution Statement

S.M.S. and J.W.R. designed experiments. S.M.S. and A.A.S. performed experiments. M.E.S. and C.A.D. analyzed the SMO microscopy results. S.M.S., M.E.S., and A.A.S. prepared figures. S.M.S., J.A.P., and J.W.R. prepared the manuscript. M.E.S., A.A.S., and C.A.D. provided manuscript feedback and edits.

## Acknowledgments

This study was funded by NIH R21 EY029167. Figure 1 is created with BioRender.com.

