## Supplementary Data for "Development and Validation of Fluorescently Labeled, Functional Type I Collagen Molecules"

### Supporting Information

#### Potential Binding Sites of Alexa Fluor 488 (AF488) on a Collagen Molecule

There are 38 positively charged and hydrophilic lysines[1] and one *N*-terminal α-amine as potential binding site on each α-chain for the label (**Figure S1**).

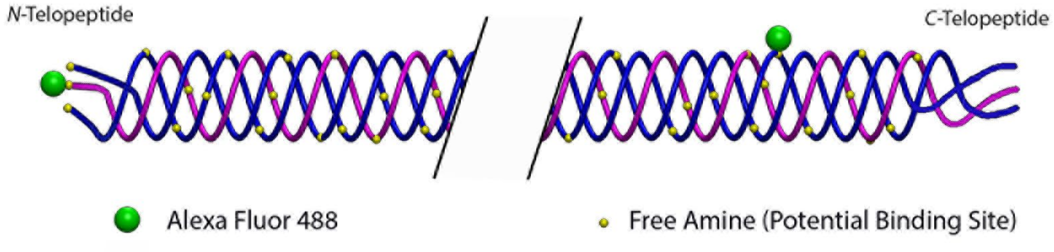


**Figure S1.** Schematic of collagen molecule labeled with 2 fluorophores. Dimensions are not to scale.

#### Effect of Labeling conditions on DOL

Monomeric collagen was labeled with AF488 and the DOL was defined as moles of dye per mole of protein. The DOL achieved at multiple pH values is shown in **Figure S2**. The initial molar ratio of dye:collagen ranged from 3-15 and the resulting dye:collagen ratio ranged from ~1 to ~10 fluorophores per collagen molecule. Tukey's honest significance test showed that increasing the initial molar ratio of AF488 to collagen increased the DOL. Furthermore, ANOVA test with a significance level of 0.05 showed that labeling performance is not significantly different at pH 7.5, 8.0, and 8.5 within each initial molar ratio of AF488:Collagen groups.

Therefore, subsequent experiments were performed at pH 7.5 to maximize the likelihood of attaching a fluorophore at the N-terminal of the monomer. It is beneficial to bind one fluorophore to N-terminus of the protein for two reasons. First, it maximizes the separation between the N-terminal bound fluorophore and a secondary fluorophore closest to the C-terminal. When the fluorophores span a greater distance on the protein, it enhances the ability to determine the orientation of the protein in SMO microscopy. Second, it has been shown that patients with Ehlers Danlos Syndrome type VII, which results in loss of the N-proteinase cleavage site in α2(I) chain, can still make collagen triple helix and have no sign of bone disease[2]. Thus, interruption of N-terminus may minimally interrupt the molecule functionality.

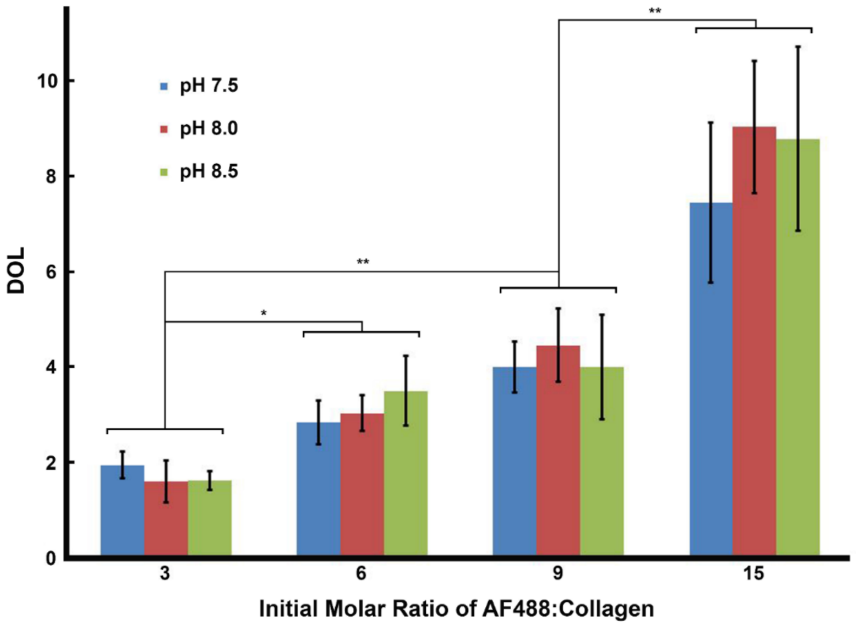


**Figure S2.** DOL achieved at pH 7.5, 8.0, and 8.5 by adding 3, 6, 9, and 15x excess moles of Alexa Fluor 488 to moles of collagen (AF488:Collagen). ANOVA test with a significance level of 0.05 showed that labeling performance is not significantly different at pH 7.5, 8.0, and 8.5 within each initial molar ratio of AF488:Collagen groups. However, labeling performance was significantly different when initial molar ratio of AF488:Collagen increased from 3 to 15 (Tukey's honest significance test; * $p<0.05$; ** $p<0.01$). Data are expressed as mean ± standard deviation (n=3 replicates).

#### De Novo Fibrillogenesis Experiments with Labeled Collagen

The kinetics of fibrillogenesis of labeled monomers was studied and compared to unlabeled monomers (DOL of 0) by measuring the lag time, plateau time, and maximum change in absorbance at 313 nm. **Table S1** summarizes the values of lag time, plateau time, and total absorbance for DOL of 0, ~0.5, ~0.8, ~1.2, ~2.1, ~4.1, and ~9.1.

**Table S1.** Absorbance, lag time, and plateau time of sigmoidal turbidity curve of 50 µg/mL labeled collagen at 37 °C. Data are presented as mean ± standard deviation, n=4 or 5 replicates per group.

| DOL | Final Absorbance (Optical Density) | Lag Time (minutes) | Plateau Time (minutes) |
|---|---|---|---|
| **0** | 0.191 ± 0.017 | 25.5 ± 0.9 | 46.7 ± 2.3 |
| **0.5** | 0.187 ± 0.016 | 25.6 ± 0.5 | 46.6 ± 1.7 |
| **0.8** | 0.182 ± 0.006 | 26.4 ± 0.8 | 46.6 ± 1.2 |
| **1.2** | 0.166 ± 0.013 | 26.8 ± 2.3 | 47.8 ± 3.8 |
| **2.1** | 0.135 ± 0.015 | 37.0 ± 1.4 | 70.8 ± 4.2 |
| **4.1** | 0.071 ± 0.015 | 34.8 ± 2.7 | 75.6 ± 7.3 |
| **9.1** | 0.051 ± 0.019 | 42.8 ± 0.8 | 67.5 ± 11.9 |

#### Proteoglycans Analysis of Scleral Fibrils

**Figure S3** shows the results of proteoglycans analysis with alcian blue. Proteoglycans concentration in the extraction solution (without collagen fibrils) and the collagen fibril solution were measured as 65 ± 4.2 and 0.25 ± 1.2 µg/mL, respectively (n = 6).

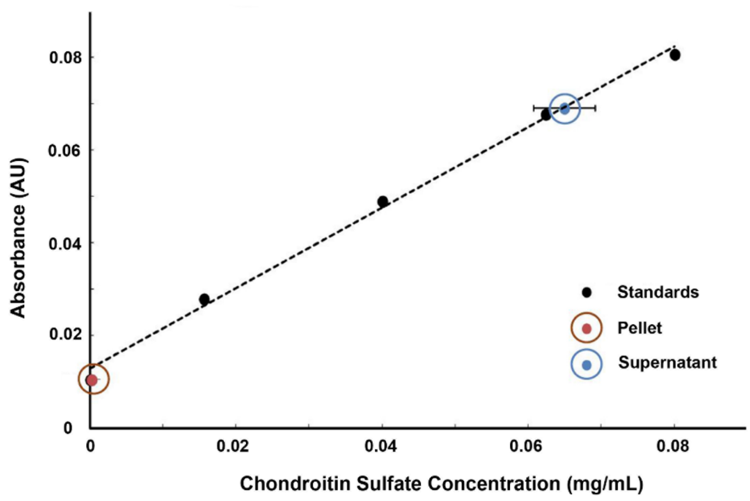


**Figure S3.** Quantification of proteoglycans with alcian blue. The concentration of proteoglycans was measured using the standard concentration of chondroitin sulfate. "Pellet" refers to the sample containing separated fibrils after extraction and "Supernatant" refers to the sample containing the fibril extraction suspension without the fibrils. Note that some error bars are too small to be seen (n = 6).

#### Fibril Diameter Measurement using TEM and DIC

Fibrils' diameter ranging between 60-240 nm can be determined using DIC if the fibril diameter distribution and the DIC edge intensity shift (DIC-EIS) distribution are known for a given batch of fibrils[3]. The diameter distribution was obtained from TEM images (**Figure S4A**). Fibrils oriented perpendicular to the shear axis of the light path (**Figure S4B**) were selected for intensity measurements to maximize signal strength and minimize error. A z-scan was used to find the maximum edge intensity shift across each fibril (**Figure S4C**).

#### AF488 Florescent Intensity Quantification

Standard samples of labeled monomers with known collagen and AF488 concentrations were used to correlate fluorescent intensity of each ROI to number of AF488 (and collagen) molecules. Fluorescent intensity of standard samples was linearly correlated to the number of AF488 molecules at our low working concentrations (**Figure S5**). Furthermore, based on the manufacturer information, AF488 is suitable for multilabeling of each protein molecule and allows for more fluorophores to be attached to each protein molecule before self-quenching becomes apparent. Therefore, no quenching between molecules was assumed.

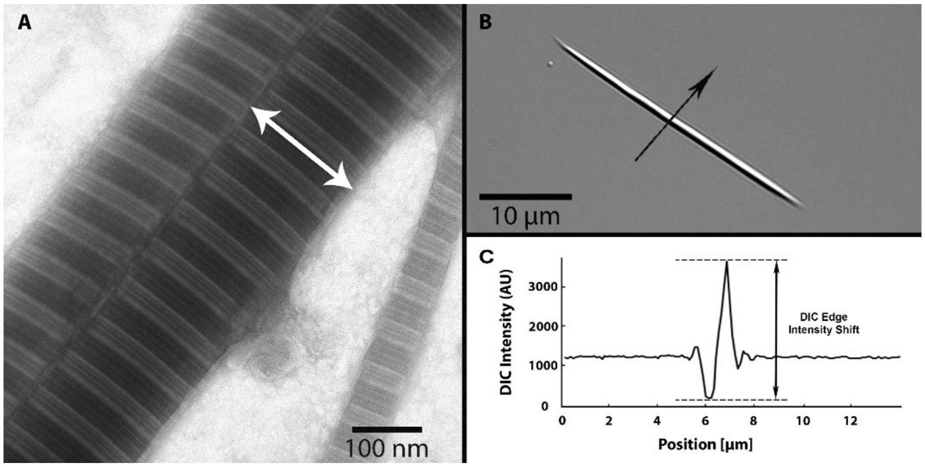


**Figure S4.** Diameter measurement of extracted sclera fibrils. (A) A typical TEM image of the trypsin extracted sclera fibrils. Images showed the native D-banding periodicity and a uniform diameter along the middle section of each fibril. (B) The DIC image shows a typical fibril (oriented in northwest-southeast direction) from the same sclera that was used for TEM imaging. (C) The intensity profile across the fibril (along the black arrow) shown in (B).

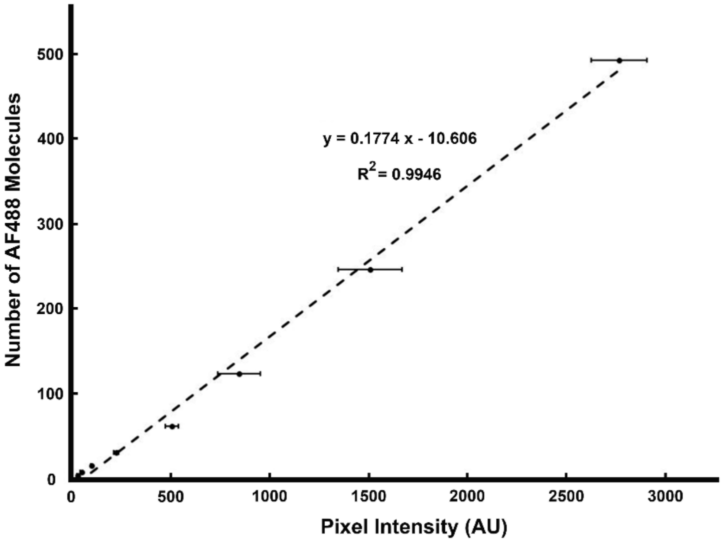


**Figure S5.** Number of AF488 molecules estimation. Fluorescent intensity of AF488 samples was linearly correlated to the number of AF488 molecules. Data points (mean ± standard deviation) represent the average pixel intensity (arbitrary unit) at known concentration of AF488 samples (n = 3).

#### **S**ingle Molecule, **M**ulti-label Fluorescence **O**rientation Microscopy (SMO Microscopy)

When the fluorescent molecules are located at two points along the collagen molecule, the epi-fluorescence image of a doubly-labeled collagen molecule is the incoherent superposition of two point-spread functions (**Figure S6A, B**) of the microscope at the emission wavelength with each one centered on one of the two fluorophores. Typically, when the separation is smaller than the resolution, the image can be approximated into an ellipse (**Figure S6C**), where the major axis represents the collagen molecule orientation. As a proof of concept, collagen fibrils were reconstituted with mixtures of labeled (DOL ~2.1) and unlabeled monomers. After reaching equilibrium, single collagen molecules that possess multiple fluorophores were detected (**Figure S6D**). The pixels of the fluorescing molecules were fitted with ellipse (**Figure S6E**), where the major axis represents the molecule orientation (**Figure S6F**). These initial results show the potential to establish a new single molecule microscopy method (SMO microscopy) capable of determining the orientation of monomers as they interact with fibrils.

We further produced synthetic images of labeled collagen monomers to estimate stochastic distribution of fluorophores along the collagen molecule and microscope parameters that will provide orientation information. Analysis of the images shows that there are subsets of the collagen molecules that generate point spread functions with equivalent major and minor axis. This can be a characteristic of a singly-labeled monomer, or a doubly-labeled monomer where the fluorophore separation exceeds the resolution of the microscope and the signal-to-noise ratio is sufficient. In the latter case, two distinct point spread functions will appear in the image, and the orientation of the fibril will be the vector direction from one to the other. However, in most cases the separation will be smaller than the resolution. In this case, a computationally simple approach can be used to calculate the covariance matrix of the two-dimensional position vector. Finding the eigenvalues and eigenvectors of this matrix approximates the image as an ellipse with the higher and lower eigenvalues representing the lengths of the major and minor axes and the corresponding eigenvectors representing their directions.

The solution of the eigenvalue problem is limited by the spacing of the fluorophores relative to the size of the point-spread function and the Poisson distribution of photons detected in each pixel of the image. Additionally, noise can result from the detector, from incomplete rejection of scattered illumination, and from autofluorescence. Although the mean of this noise can be subtracted, its variations cannot.

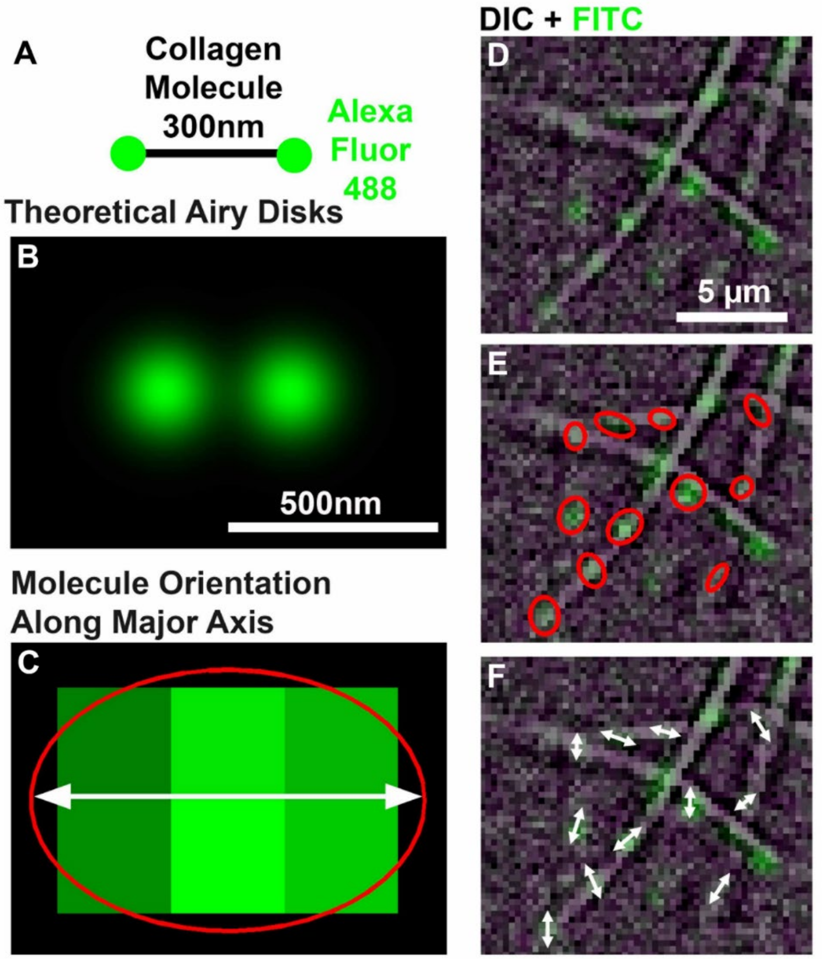


**Figure S6.** SMO microscopy showing orientation of type I bovine collagen molecules associated with native fibrils. When collagen is labeled with two fluorophores separated by a large enough distance (A), it is possible to fit an ellipse to the pixel representation (B). Thus, we can determine if the single molecules are co-aligned with the fibril axis or just associating (C, D, and E).

We have computed receiver operating characteristics (ROC) for different spacings and numbers of photons per fluorophore. To accomplish this, we generated synthetic images with Poisson random noise, computed the eccentricity as the ratio of major to minor axis and compared to a threshold. If the eccentricity exceeded threshold we recorded a detection. We repeated the simulation with one fluorophore and if the eccentricity exceeded threshold we recorded a false alarm. Each set was repeated 200 times and the probability of detection ($P_D$) was plotted against the probability of false alarm ($P_{FA}$).

Using microscope and fluorophore parameters we anticipate a maximum of a few thousand photons per fluorophore, but a low illumination level is desired in order to observe signals for a period of time without photobleaching. Thus in our computation we used between 25 and 2500 photons per fluorophore. We began with one fluorophore on each end of the monomer (spacing of 300 nm), and successively divided the spacing by 2 until the area under the ROC curves became close to 0.5.

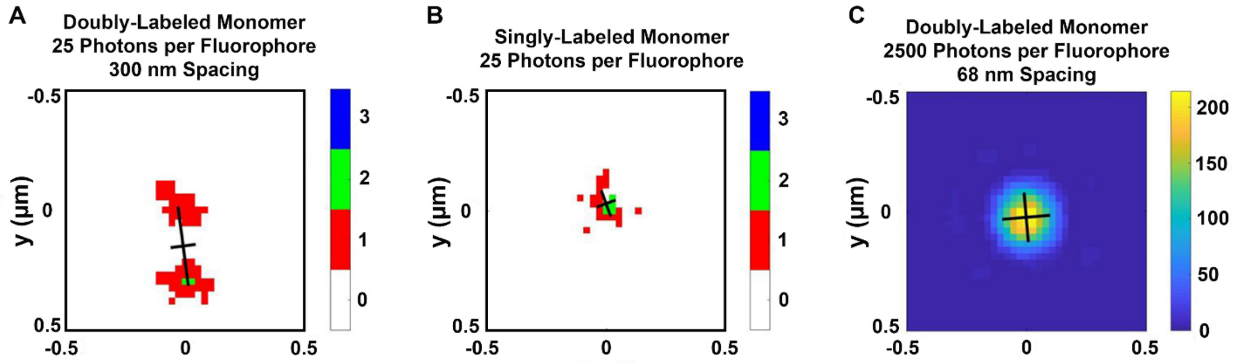


**Figure S7.** Synthetic images of labeled collagen monomer with the major and minor axes plotted. The color is representative of the number of photons per pixel. (A) Synthetic image of doubly-labeled monomer for 25 photons per fluorophore at a spacing of 300 nm. (B) Synthetic image of singly-labeled monomer for 25 photons per fluorophore for comparison. (C) Synthetic image of doubly-labeled monomer for 2500 photons per fluorophore at a spacing of 68 nm.

**Figure S7** shows three synthetic images with the major and minor axes plotted. **Figure S7A** shows the image for 25 photons per fluorophore at a spacing of 300 nm. With this large spacing, the two fluorophores are clearly resolved even with so few photons. **Figure S7B** shows a single fluorophore for comparison. **Figure S7C** shows the image for 2500 photons per fluorophore at a spacing of 68 nm. At this point it becomes very difficult to determine that there are two fluorophores present. **Figure S8** presents ROC curves for the same two spacings. The curves in each case were computed with, from top to bottom 2500, 500, 250, 100, 50, and 25 photons per fluorophore. For the larger one, the detection statistics are nearly perfect at all numbers of photons. For the smaller one, there is a slight improvement over random chance only for very large numbers of photons. We note that for low numbers of photons the ROC curve is below the 45-degree line, which represents random chance. This result occurs because with small numbers of photons, a few random detection errors can alter the symmetry of the image. The doubly-labeled monomers, with twice as many photons are less susceptible to this phenomenon.

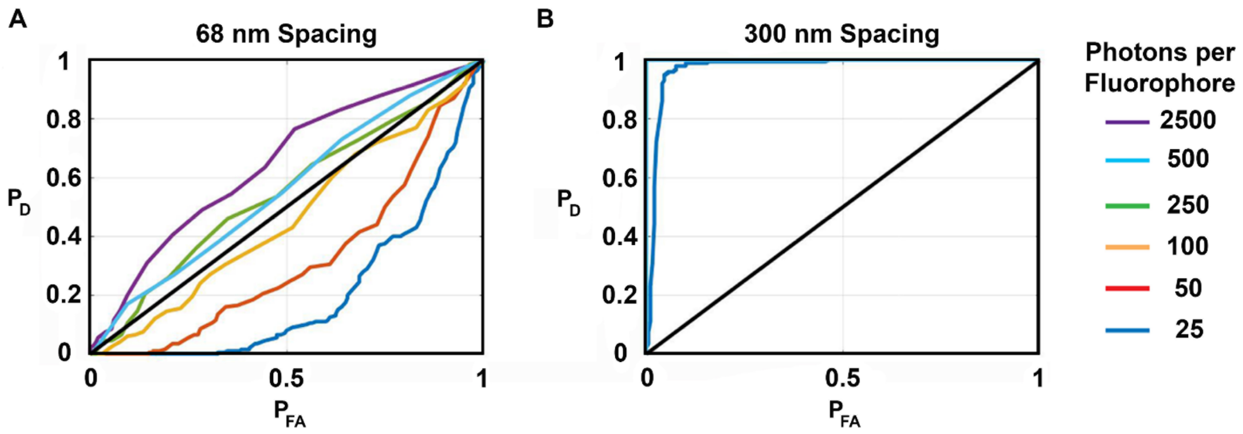


**Figure S8.** ROC curves for (A) 68 nm and (B) 300 nm spacings. The curves in each case were computed with, from top to bottom 2500, 500, 250, 100, 50, and 25 photons per fluorophore. For the 300 nm spacing, the detection statistics are nearly perfect at all numbers of photons.

Aberrations in the microscope optics, sampling limitations imposed by the camera, and non-uniform noise may further reduce performance. On the other hand, more rigorous processing approaches may improve performance. At present, we anticipate being able to determine the orientation of a useful number of monomers with fluorophore spacings exceeding about 70 nanometers. Beside the N-terminal α-amine, there are 38 lysines that are potential locations of the label. 79% of these lysines are farther than 70 nm from the N-terminal α-amine[1]. Given a high probability (e.g. more than 50%) that one of the labels is on the N-terminal, we expect that at least 40% of the labelled collagens will have detectable orientation ellipses.

Turbidity curves of 100% labeled monomers with DOL of ~2.1 shows that fibrillogenesis was affected by the labeled monomers, whereby the lag phase increased and the total absorbance decreased. Therefore, our next goal was to adjust the concentration of doubly-labelled monomers to determine what concentration ratio (labeled/unlabeled) would yield nearly normal kinetics. Our hypothesis was that including a small number of labeled monomers relative to unlabeled monomers would restore the kinetics to baseline at some threshold value. To find that threshold, we re-examined the assembly kinetics at DOL ~2.1, while the percentage of labeled monomers was varied from 100% to 1%. As shown in **Table S2**, the fibrillogenesis kinetics were not significantly altered from the unlabeled control when up to 5% labeled monomers was used. As the percentage of labeled monomers increased, the effect was more pronounced with 100% labeled monomers reducing the rate of fibrillogenesis by ~70%, increasing the lag time by ~56% and lowering the optical density by ~34%. We thus felt comfortable that using a DOL of ~2.1 which still permits tracking molecular orientation would permit us to observe the kinetics of assembly while minimizing perturbation of the process by the dye molecule.

**Table S2.** Fibrillogenesis of labeled (DOL ~2.1) and unlabeled monomers mixture. Changes in kinetics of fibrillogenesis is negligible (less than 10% error) in mixtures containing 5% or less labeled monomers. Data are presented as mean ± standard deviation (n = 3).

| Percentage of Labeled monomers | Lag Time [minutes] | Plateau Time [minutes] | Total Optical Density Change * 1000 | Normalized Slope During Growth |
|---|---|---|---|---|
| **0** | **24.8± 0.9** | **45.9 ± 2.2** | **152 ± 15** | **1.0** |
| **1%** | 24.1 ± 1.2 | 43.8 ± 2.0 | 138 ± 4 | 1.0 |
| **2%** | 22.6 ± 0.8 | 42.7 ± 2.0 | 147 ± 4 | 1.0 |
| **5%** | 22.4 ± 0.8 | 42.3 ± 2.0 | 144 ± 7 | 1.0 |
| **10%** | 26.9 ± 0.7 | 53.1 ± 4.4 | 118 ± 10 | 0.6 |
| **100%** | 36.4 ± 1.1 | 93.4 ± 20.2 | 116 ± 9 | 0.3 |

#### AF488 Absorbance Validation

AF488 concentration was calculated based on the Beer-Lambert law:

$$A = \varepsilon l c$$

where $A$ is the absorbance, $\varepsilon$ is the molar extinction coefficient, $l$ is the optical path length, and $c$ is the concentration. The labeled monomers' absorbance was measured at 494 nm in a 1 cm pathlength cuvette. Molar extinction coefficient was considered 71,000 based on the manufacturer data sheet. The results were validated using standard samples with known concentration of AF488 (**Figure S9**). Data points (mean ± standard deviation) show the measured absorbance of AF488 with known concentration (0 to $5*10^{-6}$ M). The dashed line shows calculated absorbance based on the Beer-Lambert law. The 0.033 on the dashed line equation represents the background absorbance.

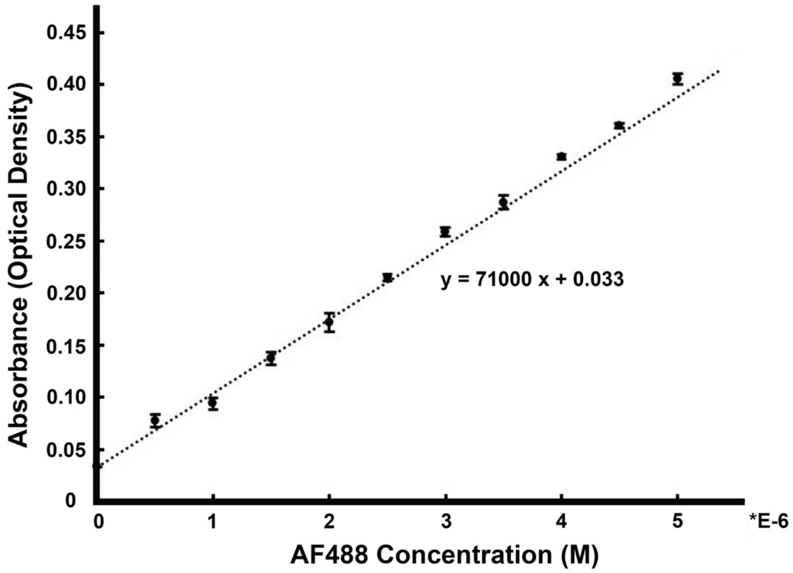


**Figure S9.** AF488 Absorbance Validation at 494 nm in a 1 cm pathlength cuvette. Data points (mean ± standard deviation) show the measured absorbance of AF488 with known concentration (n =3). The dashed line shows calculated absorbance based on the Beer-Lambert law.
